# Arrayed multicycle drug screens identify broadly acting chemical inhibitors for repurposing against SARS-CoV-2

**DOI:** 10.1101/2021.03.30.437771

**Authors:** Luca Murer, Romain Volle, Vardan Andriasyan, Nicole Meili, Liliane Yang, Daniela Sequeira, Afonso Gomez-Gonzalez, Anthony Petkidis, Dominik Olszewski, Michael Bauer, Maarit Suomalainen, Fabien Kuttler, Gerardo Turcatti, Urs F. Greber

## Abstract

Coronaviruses (CoVs) circulate in humans and animals, and expand their host range by zoonotic and anthroponotic transmissions. Endemic human CoVs, such as 229E and OC43 cause limited respiratory disease, and elicit short term anti-viral immunity favoring recurrent infections. Yet, severe acute respiratory syndrome (SARS)-CoV-2 spreads across the globe with unprecedented impact on societies and economics. The world lacks broadly effective and affordable anti-viral agents to fight the pandemic and reduce the death toll. Here, we developed an image-based multicycle replication assay for focus formation of α-coronavirus hCoV-229E-eGFP infected cells for screening with a chemical library of 5440 compounds arrayed in 384 well format. The library contained about 39% clinically used compounds, 26% in phase I, II or III clinical trials, and 34% in preclinical development. Hits were counter-selected against toxicity, and challenged with hCoV-OC43 and SARS-CoV-2 in tissue culture and human bronchial and nasal epithelial explant cultures from healthy donors. Fifty three compounds inhibited hCoV-229E-GFP, 39 of which at 50% effective concentrations (EC50) < 2μM, and were at least 2-fold separated from toxicity. Thirty nine of the 53 compounds inhibited the replication of hCoV-OC43, while SARS-CoV-2 was inhibited by 11 compounds in at least two of four tested cell lines. Six of the 11 compounds are FDA-approved, one of which is used in mouth wash formulations, and five are systemic and orally available. Here, we demonstrate that methylene blue (MB) and mycophenolic acid (MPA), two broadly available low cost compounds, strongly inhibited shedding of infectious SARS-CoV-2 at the apical side of the cultures, in either pre- or post-exposure regimens, with somewhat weaker effects on viral RNA release indicated by RT-qPCR measurements. Our study illustrates the power of full cycle screens in repurposing clinical compounds against SARS-CoV-2. Importantly, both MB and MPA reportedly act as immunosuppressants, making them interesting candidates to counteract the cytokine storms affecting COVID-19 patients.

## Introduction

Coronaviruses (CoVs) are enveloped single-strand, plus-sense RNA viruses. Based on their genomic sequence and phylogenetic relationship, the *Coronavirinae* subfamily is divided into four genera, alpha-, beta-, gamma- and delta-CoV (1). In December 2019, a local outbreak of pneumonia caused by a previously unknown CoV was reported in Wuhan (Hubei, China). The causative agent was identified as 2019-nCoV, sequenced and found to be a beta-coronavirus (2). It was later named as SARS-CoV-2. SARS-CoV-2 is the causative agent of COVID-19, a pandemic disease, which has caused more than 126 million PCR positive cases and 2.7 million deaths (status March 28, 2021, computed by the Johns Hopkins Coronavirus Research Center: <u>(3)</u>.

Severe cases of COVID-19 are accompanied by respiratory failure, and require mechanical ventilation at intensive care units (ICU). Patients with severe COVID-19 are at risk to develop acute respiratory distress syndromes (ARDS), multiple organ dysfunction syndromes (MODS) or failures (MOF) and death {32150360}. Current treatment options for patients with severe COVID-19 are limited, and include supportive care, administration of the viral RNA polymerase inhibitor remdesivir together with corticosteroids to limit inflammatory distress in COVID-19 (https://www.covid19treatmentguide-lines.nih.gov/therapeutic-management/). As SARS-CoV-2 continues to adapt to humans and evades immune pressure, vaccinations as well as antibody therapies may be insufficient treatment options in the longer run to contain the disease (4,5). Additional treatments, including low cost chemical compounds with sufficient efficacy and safety are urgently needed.

Drug repurposing allows for rapid identification of clinically approved or investigational compounds towards emerging indications. This approach offers a multitude of advantages over *de novo* drug development (reviewed in (6)). Most importantly, candidate compounds have a sufficient safety record under well defined conditions, allowing for direct transition to human clinical trials. At this critical stage of the COVID-19 pandemic, drug repurposing can massively increase the treatment options for COVID-19 cases with bad prognoses. Several drugs have already been proposed for repurposing against COVID-19, including antivirals, such as remdesivir at high cost, or low cost anti-malaria compounds, such as chloroquine (7), for which clinical trials were suspended due to inefficacy (8). Investigational studies identified highly promising drug candidates, some of which are now in clinical trials, such as the translation elongation inhibitor Aplidin approved for the treatment of multiple myeloma (9–11). A co-formulation of Aplidin plus dexamethasone is currently in phase III clinical trials against COVID-19 (NCT04784559).

Here, we present the results from a multicycle drug repurposing screen which uncovers drug candidates for COVID-19 treatment. Unlike canonical drug screens, our assay sampled the entire replication cycle of a GFP expressing variant of the endemic α-coronavirus hCoV-229E in the human hepatoma cell line Huh7. We validated the initial hits by two additional hCoVs, OC43 and SARS-CoV-2 in nasal and bronchial human airway epithelial explant cultures (HAEEC) grown at air-liquid interface, and identified 11 compounds with broad coronavirus effects, one topical compound (Cetylpyridinium chloride monohydrate) used in mouth wash formulations (12), and 10 systemically used compounds. Five of them are approved for human use, namely methylene blue (MB), mycophenolic acid (MPA), Posaconazole, Thonzonium bromide and R788-Fostamatinib. Two systemic compounds are in phase II clinical trials (MLN4924, Ravuconazole), and three in preclinical tests (GPP-78, Ro 48-8071, SB-505124). Interestingly, two of the approved compounds have combined anti-viral and reportedly anti-inflammatory effects. They are broadly available at low cost.

## Results

### Compound identification by image-based full infection-cycle screening with hCoV-229E-GFP

Host targeting with chemical compounds combined with full cycle image-based analyses is a powerful approach against viral infections, as recently demonstrated with the identification of the HIV protease inhibitor Nelfinavir blocking human adenovirus egress from infected cells (13,14). Here we adapted a similar approach to identify broadly effective coronavirus inhibitors. The overall procedure of our anti-coronavirus compound screen is depicted in Fig. 1. Starting with a chemical library of 5440 mostly repurposable compounds we screened for inhibition of hCoV-229E-GFP plaque formation in an arrayed 384 format using Huh7 cells and remdesivir as a positive control. Z’ factors (computed as described in (15)) were mostly at or above 0.5, entailing a high robustness, excellent assay quality and consistent analysis pipelines (Fig. S1). The screen yielded 53 hits, which were validated for effective concentrations (EC_50_) and toxicity (TC_50_) in hCoV-229E-GFP infected Huh7 cells. Two subsequent validation filters were applied with hCoV-OC43 infection of Huh7 cells, and SARS-CoV-2 infection of VeroE6, Huh7 expressing the angiotensin-converting enzyme 2 (ACE2), HeLa-ACE2 and A549-ACE2 cell lines. A set of selected drugs was then tested in nasal and bronchial HAEEC. These procedures identified MPA and MB to be effective against SARS-CoV-2 infection, and in particular inhibiting virus cell-cell transmission.

**Figure 1:**
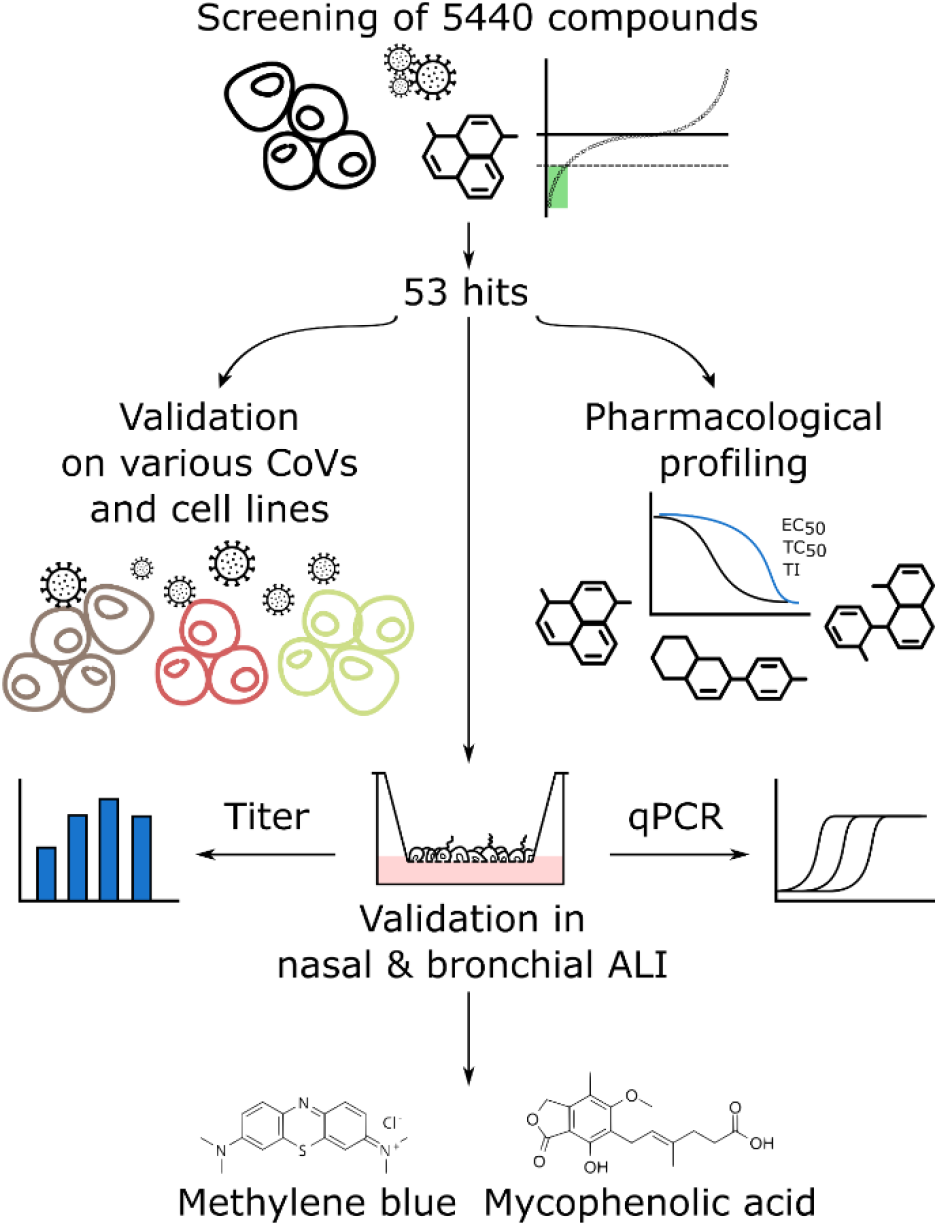
Overall workflow of the screen. Initially, we assessed a library of 5440 compounds for efficacy against hCoV-229E-GFP. This yielded 53 hits. We then tested these hits for efficacy against hCoV-OC43 and SARS-CoV-2 in different cell lines, including Vero, Huh7-ACE2, A549-ACE2 and HeLa-ACE2. In parallel, we determined the EC_50_, TC_50_ and the ratio TC_50_ / EC_50_, i.e., the putative therapeutic index (TI). A selection of hits in advanced clinical state (approved and systemically administrable) was tested for anti-SARS-CoV-2 inhibition in differentiated nasal and bronchial airway epithelia grown at air-liquid interface. Methylene blue (MB) and mycophenolic acid (MPA) were the most potent inhibitors of infectious SARS-CoV-2 particle formation.

We employed the EPFL BSF-repurposing compound collection, curated and corrected based on information made available by the Broad Institute (16). The library comprises 39% clinical compounds, 26% are in phase I – III trials, 34% in preclinical development and 1% have been withdrawn. The collection covers substantial chemical space with 5032 clusters, 4641 of which contain a single compound, as determined by sphere exclusion clustering analysis with a Tanimoto distance of 0.3 between centroids. One thousand two hundred eighty compounds were contained within the Prestwick Chemical Library (PCL) and were purchased as such. The remaining 4160 were purchased from 34 different suppliers, including MedChem Express (59.5%), MolPort (35.5%) and others. All compounds were interrogated for chemical integrity and purity by LC-MS, and were assessed for toxicity by PrestoBlue staining (Fig. S2), a resazurin based assay for measuring overall energy levels of cell populations (17). These results are in good agreement with the number of segmented nuclei based on Hoechst 33342 staining of the screening plates (see Table S1). The library was arrayed in microscopy grade 384-well assay plates, followed by cell seeding, infection and high-throughput imaging, as described (14,18) (Fig. 2A). This allowed us to extract image analysis parameters, including cell count, infected cell count, total GFP intensity and number of infection foci. These parameters were normalized perplate to the mean value of all negative control wells. Compounds were classified as hits inhibiting infection foci (plaque) formation if a Z score cutoff of -3 for any of these normalized infection parameters was obtained (Fig. 2B). Compounds were flagged toxic if the mean cell count as determined by nuclear counterstaining was below the mean of all negative control wells minus 2 standard deviations of all negative control wells. Analysis and post-processing yielded a total of 53 compounds fulfilling the criteria described above. The hit compounds were classified either as antiseptics, antifungals, antibacterials, anticancer, metabolism as well as anti-inflammatory agents (Fig. 2C). Among the immuno-suppressant and anti-inflammatory compounds, we selected MB and MPA for further validations. MB effectively reduced the number of infection foci. MPA reduced the total GFP intensity, while remdesivir showed complete infection inhibition of hCoV-229E-GFP at 0.33 µM (Fig. 2D).

**Figure 2:**
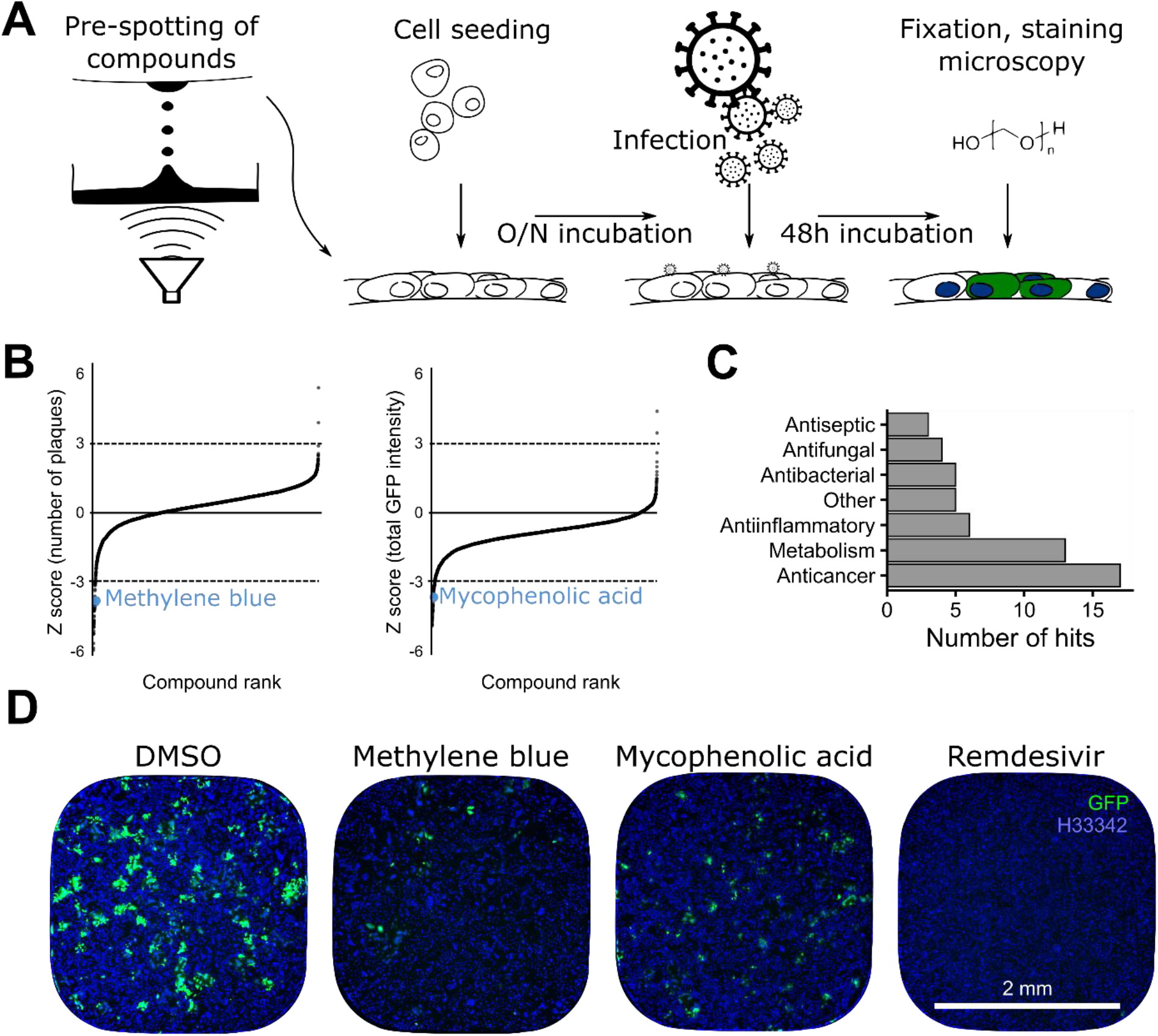
High level view of the compound library screen. **A)** Schematic representation of the screening procedure. **B)** Compound ranking by number of plaques and total GFP intensity. The cut-off value was set at a Z score of -3. MB and MPA are highlighted in blue. **C)** Most of the hits classify as anticancer agents or modulators of metabolism. **D)** Example images of wells treated with the indicated compounds. In the negative control (DMSO), clusters of GFP positive cells represent infection foci (green). Nuclei are represented in blue.

### Dose-response validation and hit extension to hCoV-OC43 and SARS-CoV-2

To assess the ther-apeutic potential of the 53 hit compounds EC_50_ and TC_50_ values were determined for hCoV-229E-GFP in Huh7 cells by conducting dose-response experiments with concentrations from 1.525 nM to 50 μM. The results are summarized in Table 1 and Table S1. Thirty nine compounds had an EC_50_ < 2μM and a therapeutic index (TI) of > 2, as determined by fluorescent focus formation (FFF). The slight discrepancy to the results obtained in the initial screen at 1.67 µM was due to additional parameters used for hit scoring in the original screen, including number of GFP positive cells, total GFP intensity and infection index, in addition to the number of fluorescent foci. Notably, the dose-response analyses yielded LY2090314 as the most potent compound, where the lowest concentration tested (1.525 nM) resulted in a 41% reduction of infection foci. We conservatively estimated of an EC_50_ of about 2 nM, and a TI of 15830 based on an observed TC_50_ of 31.7 μM. MB had an EC_50_ of 1.43 μM, a TC_50_ of 8.71 μM (TI of 6.09), and MPA EC_50_ was estimated to 1.85 μM at a TC_50_ of 45.8 μM, resulting in a TI of 24.8 (Fig. 3A). In addition, 39 compounds used at 1.75 μM proved to be effective and non-toxic against hCoV-OC43 infection of Huh7 cells (Table 1).

**Table 1:** Validation of the 53 top hits. All compounds qualifying as hits in the original screen are listed. The compounds highlighted in bold at the top of the table are clinically launched and in clinical use. Columns represent different validation experiments with the corresponding viruses and cell lines. Ticks mark compound qualifying as a hit in the respective experiment. There is a good agreement between the original screen, 229E-GFP dose-response (D-R) and hCoV-OC43 on Huh7 with fewer compounds also being effective against SARS-CoV-2.

| Compound | 229E-GFP<br>D-R Huh7 | OC43<br>Huh7 | SARS-CoV-2<br>Huh7-ACE2 | SARS-CoV-2<br>VeroE6 | SARS-CoV-2<br>HeLa-ACE2 | SARS-CoV-2<br>A549-ACE2 |
| --- | --- | --- | --- | --- | --- | --- |
| **Cetylpyridinium** | ✓ |  |  |  | ✓ | ✓ |
| **Fostamatinib** | ✓ | ✓ | ✓ |  |  | ✓ |
| **Methylene Blue** | ✓ |  |  |  | ✓ | ✓ |
| **Mycophenolic acid** | ✓ | ✓ | ✓ |  | ✓ | ✓ |
| **Posaconazole** |  | ✓ | ✓ | ✓ | ✓ | ✓ |
| **Thonzonium bromide** | ✓ | ✓ | ✓ |  | ✓ | ✓ |
| A0001 | ✓ |  |  |  |  |  |
| Abafungin | ✓ | ✓ |  |  | ✓ |  |
| Acivicin | ✓ | ✓ |  |  |  |  |
| Adarotene |  | ✓ |  |  |  |  |
| Amuvatinib | ✓ | ✓ |  |  |  |  |
| Anisomycin | ✓ | ✓ |  |  |  |  |
| APY0201 |  | ✓ |  | ✓ |  |  |
| AT9283 | ✓ | ✓ |  |  |  |  |
| AZD-5438 | ✓ | ✓ |  |  |  |  |
| AZD2858 | ✓ | ✓ |  | ✓ |  |  |
| Betulinic acid | ✓ | ✓ |  |  |  |  |
| Cerdulatinib | ✓ | ✓ |  |  |  |  |
| Cetalkonium chloride | ✓ | ✓ |  |  | ✓ |  |
| CHIR-124 | ✓ | ✓ |  |  |  | ✓ |
| CHIR-98014 | ✓ | ✓ |  |  |  |  |
| CUDC-907 |  |  |  |  |  |  |
| Cyclopiazonic-acid | ✓ | ✓ |  |  |  |  |
| Eprinomectin | ✓ |  |  |  |  |  |
| FH535 | ✓ |  |  |  |  |  |
| Geldanamycin |  | ✓ |  |  |  |  |
| GPP-78 | ✓ | ✓ | ✓ |  |  | ✓ |
| GSK 3 Inhibitor IX | ✓ | ✓ |  |  |  | ✓ |
| GZD824 | ✓ | ✓ |  |  |  | ✓ |
| Isavuconazole | ✓ |  |  |  |  |  |
| JTE-013 | ✓ | ✓ |  |  |  |  |
| LE-135 | ✓ | ✓ |  |  |  |  |
| Lomefloxacin HCl |  |  |  |  |  |  |
| LY2090314 |  | ✓ |  | ✓ |  |  |
| Mebendazole |  | ✓ |  | ✓ |  |  |
| MLN4924 | ✓ | ✓ |  | ✓ | ✓ | ✓ |
| OTSSP167 | ✓ |  |  |  |  |  |
| Oxaprozin |  |  |  |  |  |  |
| Pelitinib | ✓ | ✓ |  |  |  | ✓ |
| PF-04457845 | ✓ | ✓ |  |  |  |  |
| PF-3845 | ✓ | ✓ |  |  |  |  |
| PFK-015 |  | ✓ |  |  |  |  |
| RAF265 | ✓ | ✓ |  |  |  | ✓ |
| Ravuconazole | ✓ | ✓ | ✓ |  |  | ✓ |
| Ro 48-8071 | ✓ | ✓ | ✓ | ✓ | ✓ | ✓ |
| SB-505124 |  |  |  |  | ✓ | ✓ |
| SB225002 |  | ✓ |  |  |  |  |
| Tanaproget | ✓ | ✓ |  |  |  |  |
| TCS-21311 | ✓ | ✓ |  | ✓ |  |  |
| TRO 19622 |  |  |  |  |  |  |
| UK-5099 | ✓ | ✓ |  |  |  |  |
| Verteporfin | ✓ |  |  | ✓ |  |  |
| VU29 |  |  |  |  |  |  |
| Total | 39 | 39 | 7 | 9 | 10 | 16 |

**Figure 3:**
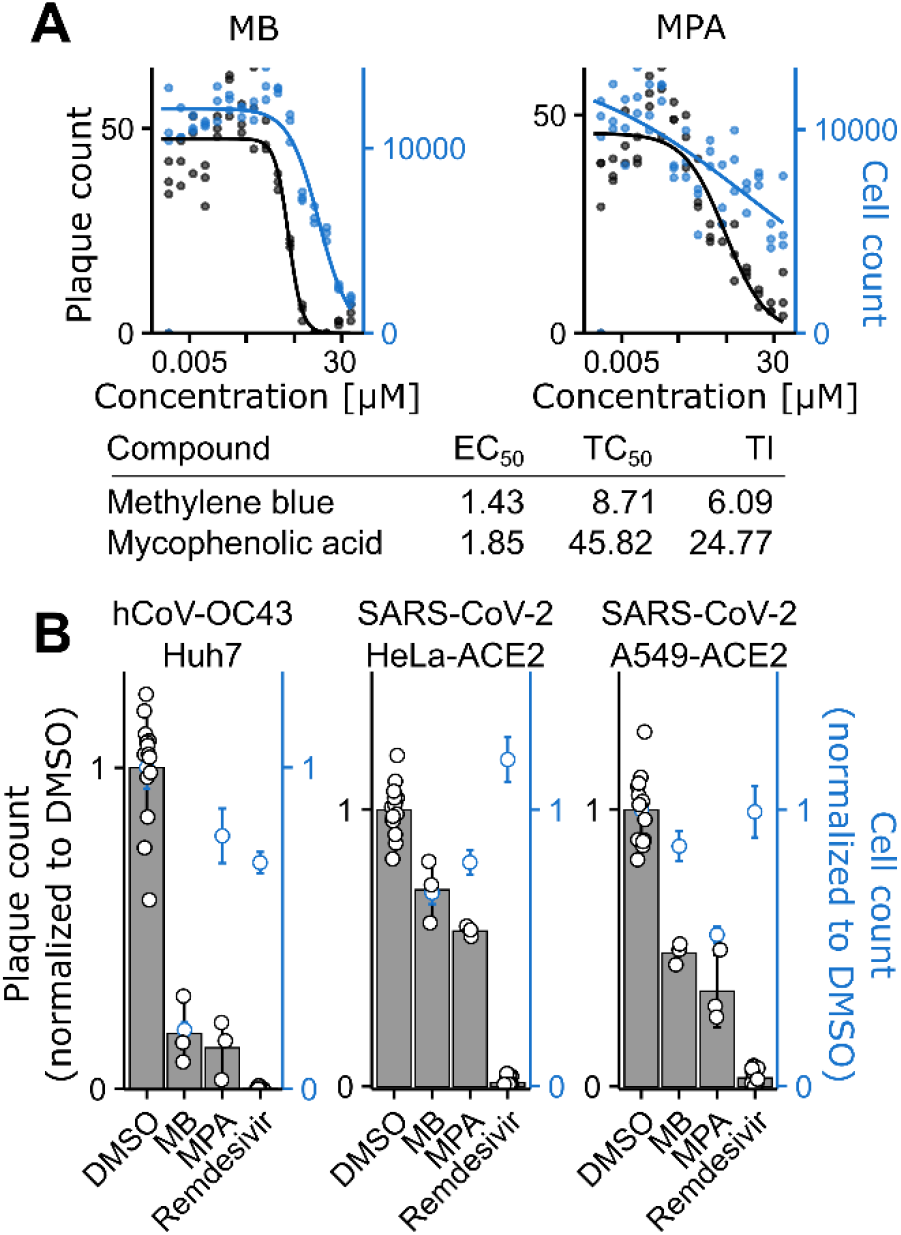
Validation of MB and MPA. **A)** Dose response curves with MB and MPA treated Huh7 cells infected with hCoV-229E-GFP at 50 FFU in presence of the indicated concentration of compound. After 48h, cells were fixed and imaged. Curves were fitted through the data points using a 3-parameter log-logistic function with a lower limit of 0. The extracted values are summarized in a table below the graphs. **B)** Indicated cell lines were infected with either hCoV-OC43 or SARS-CoV-2 in presence of MB or MPA for 24h or 48h. Cells were stained for viral nucleoprotein production. Both MB and MPA effectively inhibited viral plaque formation.

We next generated Huh7, human lung epithelial A549 and human cervical carcinoma HeLa cells expressing ACE2 by lentivirus transduction (Fig. S3). These ACE2 transduced cells as well as African green monkey kidney epithelial VeroE6 cells were susceptible to SARS-CoV-2 (Fig. 3B and Table 1). Eleven compounds were effective in two or more cell lines against SARS-CoV-2, and two of them in all cell lines, namely Posaconazole, and Ro 48-8071. The preclinical compound Ro 48-8071 had the best hit profile regarding effectiveness in all four cell lines against SARS-CoV-2, and hCoV-OC43. Ro 48-8071 affects oxidosqualene cyclase in cholesterol bio-synthesis {Maione, 2015, 25761781}. In addition, the launched anti-fungal compound Posaconazole showed a similar profile, but did not meet the dose-response criterion with an EC_50_ of 4.20 μM. We did, however, not observe any toxicity effects for Posaconazole up to 50 μM, indicating it is well tolerated by cells. MPA, thonzonium bromide and MLN4924 were effective against SARS-CoV-2 in three cell lines, and also inhibited hCoV-OC43. Notably, the number of compounds that were effective against both hCoV-229E-GFP and hCoV-OC43 was much higher than those effective against SARS-CoV-2. This effect did not seem to be cell-line dependent, as for example Huh7-ACE2 used for SARS-CoV-2 are derived from Huh7 cells that were used for hCoV-229E-GFP and hCoV-229E. Possibly though, different temperatures used for SARS-CoV-2 and 229E or OC43 infections (37°C versus 33.5°C) may affect the effectiveness of some of the drugs against SARS-CoV-2. In summary, the cell line screens with the endemic coronaviruses 229E, OC43 and the pandemic SARS-CoV-2 uncovered five clinically launched compounds, MB, MPA, Posaconazole, Thonzonium bromide, and R788-Fostamatinib plus the mouth wash compound Cetylpyridinium (Table 1).

### MB and MPA inhibit SARS-CoV-2 infection of human airway epithelial explant cultures

Based on the above broad antiviral effects in cell cultures, we next selected three of the five systemically available launched compounds, plus four promising preclinical compounds for anti-SARS-CoV-2 assessment in human nasal and bronchial HAEEC, sold under the brand name MucilAir™ (Table 2). The launched compounds comprised MB, MPA, and R788-Fostamatinib, but not Posaconazole and Thonzonium bromide, which are subject to another study. None of the four preclinical compounds tested, the VEGF receptor and Raf1 inhibitor RAF265 (19), the Bcr-Abl / Src inhibitor GZD824 (20), the vitamin E metabolite A0001 (a-tocopheryl-quinone) (21), and the broad kinase inhibitor OTSSP167 (22) showed anti-viral effects distinct from toxicity.

**Table 2:** Compounds tested on HAEEC. The listed compounds were tested for efficacy against SARS-CoV-2 in human differentiated nasal and bronchial airway explant cell cultures. The compounds with variety of original indications are generally in advanced clinical trials or approved. The final column denotes the number of cell lines out of 4, which showed inhibition of SARS-CoV-2 with a given compound.

| Compound | Original indication | Clinical status | EC<sub>50</sub> [μM] | TC<sub>50</sub> [μM] | Susceptible cell lines (n.) |
| --- | --- | --- | --- | --- | --- |
| A0001 | Friedreich ataxia | Phase II | 0.497 | 8.4 | 0 |
| Fostamatinib | Immune thrombocytopenic purpura | Launched | 1.331 | 14.709 | 2 |
| GZD824 | Cancer | Experimental | 0.49 | 0.98 | 1 |
| Methylene Blue | Methemoglobinemia | Launched | 1.426 | 8.709 | 2 |
| Mycophenolic acid | Immunosuppressant | Launched | 1.85 | 45.823 | 3 |
| OTSSP167 | Cancer | Experimental | 0.041 | 0.766 | 0 |
| RAF265 | Cancer | Phase II | 0.698 | 6.963 | 1 |

Well differentiated human nasal or bronchial airway cells grown at air-liquid interface were either pre-treated with low micromolar basolateral compounds for 2 h, or treated with compounds at 1 day post inoculation with SARS-CoV-2 to the apical (airway) exposed side. All three clinically launched drugs were well tolerated in the HAEEC for at least 18 d with periodic addition of fresh compound to the basolateral medium (not shown). While both MB and MPA were effective at inhibiting SARS-CoV-2 infection, R788-Fostamatinib exhibited no measurable effects on SARS-CoV-2 titer release in the apical milieu at 3.3 or 10 µM, in contrast to remdesivir, which was effective at 10 µM in both pre- and post exposure settings (Table 2, and Fig. 4A, B), the latter as previously reported (23).

**Figure 4:**
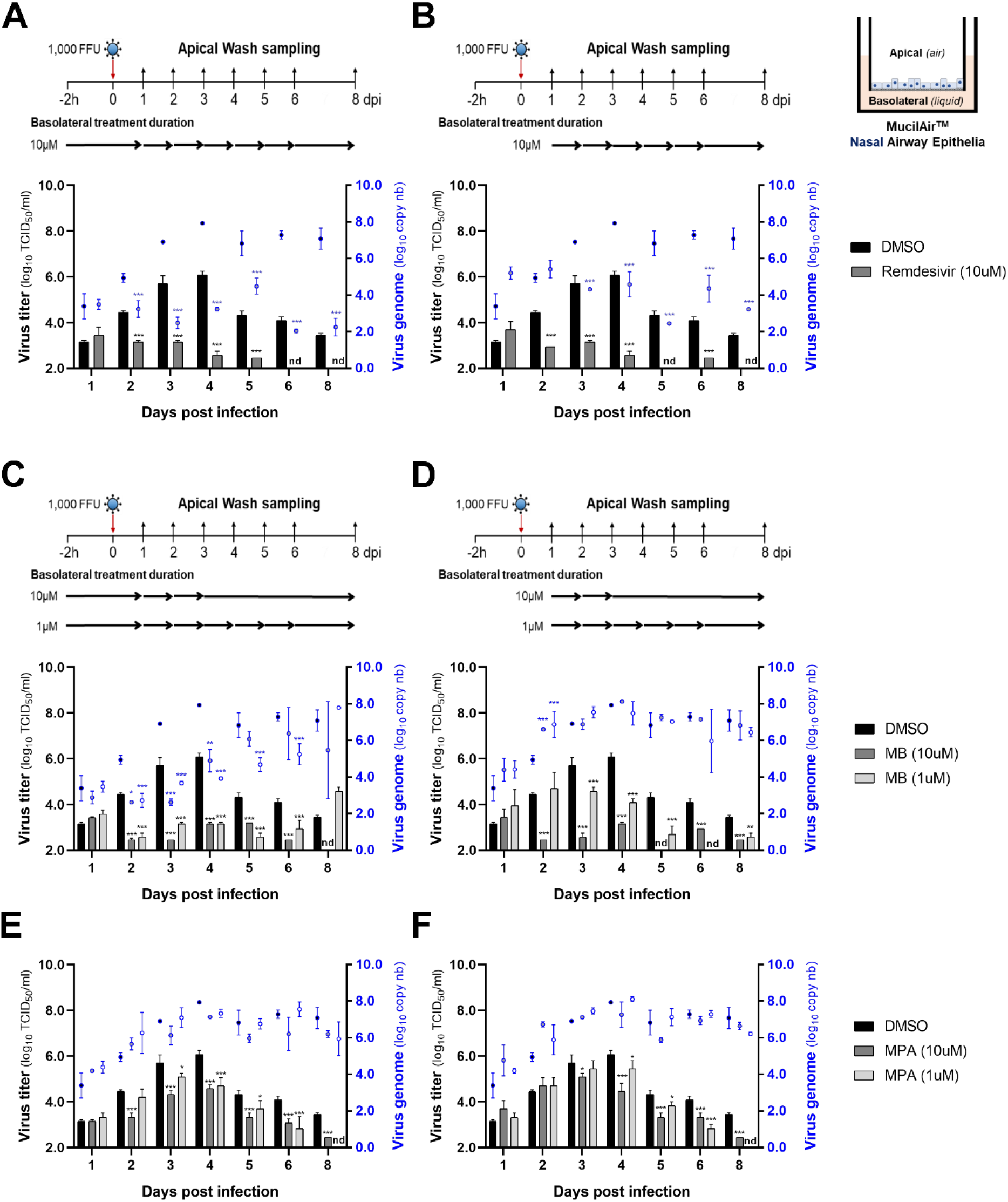
Inhibition of SARS-CoV-2 infection of nasal HAEEC by MB and MPA. HAEEC (MucilAir, Epithelix) grown at air-liquid interface were inoculated apically with 1,000 FFU of SARS-CoV2 (day 0), and subjected to drug treatment in the basolateral medium. Drugs were applied in a pre-infection regimen, starting at 2h prior to SARS-CoV2 inoculation (**A, C, and E**) or in a post-infection manner starting at 1d post inoculation (**B, D, and F**). Remdesivir (10µM) served as a reference. Compounds were daily administrated until d8 (**A-B**). MB and MPA were administrated daily at 10 and 1 µM until day-3 and day-6 post infection, respectively, then in daily mode until day-8 (**C-F**). SARS-CoV2 produced at the apical side was collected daily by apical washing and quantified by virus TCID_50_ titration (*bars, y-axis on the left*), and by RT-qPCR of the SARS-CoV2 membrane protein gene (*blue dots, y-axis on the right*). Data represent the mean±SD of two independent replicates. Statistical significance (p-value) was calculated by multiple t-tests; p<0.05 (*), p<0.01 (**), p<0.001 (***) versus the DMSO control. Not determined (nd) indicates virus titers below the titration level of 2.4 log_10_ TCID_50_/ml.

In contrast to Fostamatinib, both MB and MPA treated cells remained morphologically unchanged, even in presence of virus. MB and MPA treated cells displayed robust ciliary beating akin to uninfected cells, but distinct from infected cells in absence of compound which showed decreased ciliary motility as indicated 6 dpi (Suppl. Movies 1-5). The treatment with MB at 1 or 10 µM was very effective and blocked the release of infectious virus particles by up to 2 logs at 4 dpi and to near undetectable levels 8 dpi, while DMSO-treated cells released massive virus titers peaking at 6.1±0.2 log_10_ TCID_50_/ml (Fig. 4C, D). The strong antiviral effect of MB was reflected in a reduction of viral genome load in the apical milieu by about 2 logs until 3-4 dpi, as indicated by RT-qPCR measurements. The rt-qPCR measurements were internally controlled by linear regression analyses of the Ct values from RNA standards against their respective copy numbers in the range of 10^3^ to 10^6^ per µl yielding R^2^ correlation coefficients of 0.9657 with inter-assay (n=8) coefficients of variation ranging from 3.2 to 3.8% (Fig. S4). In addition, our PCR measurements gave comparable results with probes for either the M or the Pol gene (Nsp12), the latter only present in genomic but not subgenomic RNA (24), thereby ruling out the possibility that subgenomic RNA fragments were preferentially released from the infected cells (Table S2 and S3). Intriguingly, at 5-8 dpi, the apical genome copies were no longer significantly reduced by MB, although the viral titers were reduced to near undetectable levels. Likewise, MPA had significant anti-viral effects, albeit less pronounced than MB, and with delayed post-exposure efficacy particularly at 10 µM (Fig. 4E, F). Intriguingly, the genome copy numbers in the apical supernatant were only mildly reduced by MPA. Nevertheless, the anti-viral effects of both MB and MPA were reproduced in bronchial HAEEC treated with either 0.5 or 5 µM of MB or 1 or 10 µM MPA (Fig. 5A-D).

**Figure 5:**
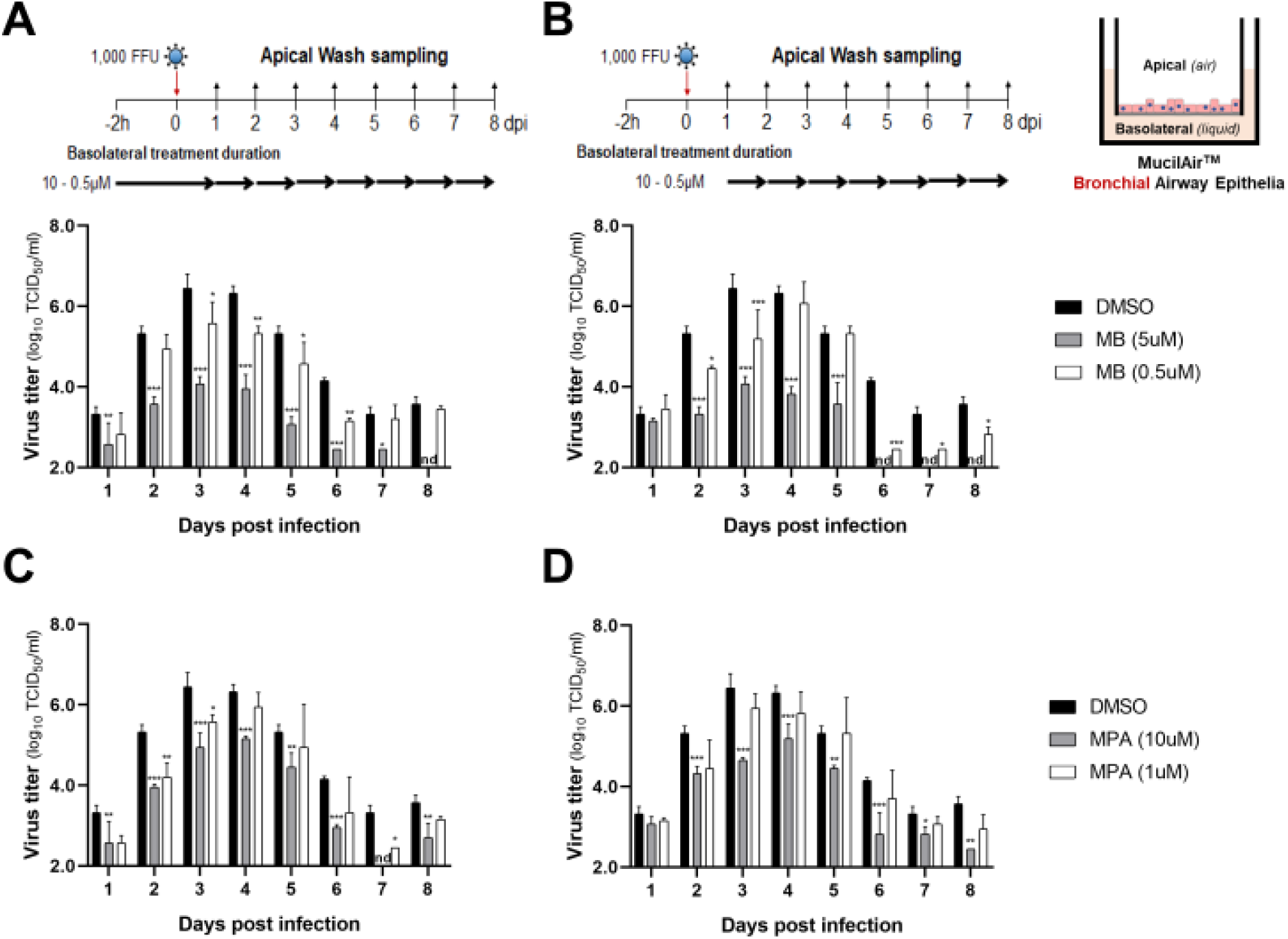
Inhibition of SARS-CoV-2 infection of bronchial HAEEC by MB and MPA. HAEEC grown at air-liquid interface were inoculated apically with 1,000 FFU of SARS-CoV2 (day 0), and subjected to drug treatment in the basolateral medium. Drugs were applied in a pre-infection regimen, starting at 2h prior to SARS-CoV2 inoculation (**A, C**) or in a post-infection manner starting at 1d post inoculation (**B, D**), and MPA at 10 and 1 µM until day-8 post infection (**C-D**). DMSO was used as a negative control. SARS-CoV-2 released to the apical side was collected daily by apical washing and quantified by virus TCID_50_ titration (*bars, y-axis on the left*). Bars with whiskers represent the mean±SD of two independent replicates. Statistical significance (p-value) was calculated by multiple t-tests; p<0.05 (*), p<0.01 (**), p<0.001 (***) versus the DMSO treated control. Not determined (nd) indicates virus titers below 2.4 log_10_ TCID_50_/ml.

## Discussion

SARS-CoV-2 and its genetic variants spread across the globe with unprecedented impact. Although vaccinations are ramped up on a global scale, it will take years to deliver the vaccine to all those who wish to receive it. In addition, individuals suffering from COVID-19 are in need of acute medical treatment, and would greatly benefit from broadly available safe and effective anti-virals with post exposure efficacy. Likewise, given the zoonotic and anthroponotic transmission of coronaviruses from animals to humans and from human to animals (25,26), the treatment of livestock and pets with compounds effective against SARS-CoV-2 may be considered in the future to restrict viral reservoirs. Accordingly, host targeting is likely less prone to raise viral resistance, unlike direct targeting of the virus, for example by remdesivir or the mutator nucleoside MK-4482/EIDD-2821 (27–30). Notably, remdesivir leads to stalling of the coronavirus RNA-dependent RNA polymerase and has a conserved mode of inhibition across different viruses, where a single point mutation in the Ebolavirus polymerase mediates drug resistance (29,31).

*De novo* development of antivirals is a lengthy process at high cost and impracticable to resolve the current crisis in short terms {Ward, 2015, 26576812}. Our large scale drug repurposing screen identified broadly acting clinically approved small molecular weight compounds. We took advantage of EPFL’s curated and mass spectrometry-validated BSF library composed of 5440 distinct chemical compounds, most of which have been approved for human use in nonviral indications or in clinical development. We harnessed the power of fluorescence imaging assessing the full virus replication cycle from entry to egress and spread to neighboring cells. This procedure previously identified Nelfinavir as an adenovirus egress inhibitor, notably in contrast to a previous publication earmarking it as ineffective against adenovirus in PCR-based virus replication assays (14,32).

The coronavirus screen developed here scored fluorescent focus formation of hCoV-229E-GFP, validated the hits by toxicity assessments, counter screened with a second endemic coronavirus hCoV-OC43, and finally validated the potency of the compound hits by SARS-CoV-2 infection. It is unlikely, though not impossible, that our full cycle screen discovers coronavirus entry inhibitors, since the entry pathways of endemic hCoVs are distinct from SARS-CoV-2 (25). For example, the hCoV-229E S-glycoprotein binds to aminopeptidase N, hCoV-NL63 lacking a furin cleavage site binds to the ACE2 receptor, HKU1 and OC43 bind to O-acetylated sialic acid (33– 35). SARS-CoV and SARS-CoV-2 bind to ACE2 (36). Upon furin cleavage of the S-glycoprotein, the latter contacts neuropilin for efficient entry (37,38). Hence, the compounds that we identified here likely target host functions required for virus progeny formation, egress from infected cells or transmission to neighboring cells.

### Clinical agents for repurposing against COVID-19

The top clinical agents from our investigation are MB and MPA. Both scored as hits with the endemic hCoV as well as SARS-CoV-2 cell culture infections and against SARS-CoV-2 in nasal and bronchial HAEEC. MB was previously proposed for COVID-19 treatment based on its inhibition of SARS-CoV-2 replication in Vero-E6 cells (39). The compound is broadly used in humans, for example in peroral formulation against malaria (40,41), or in anti-methemoglobinema treatment (42). Many trials use it as a placebo control making it one of the best controlled compounds in clinics. However, MB is not inert. It is a redox-cycling agent switching between the oxidated form (MB) and the reduced form, called leuco-MB (L-MB) (43). The reduction occurs through cellular flavo oxido-reductases, such as glutathione reductase, thioredoxin reductase or the mitochondrial membrane associated dihydrolipoamide dehydrogenase at the expense of nicotinamide adenine dinucleotide phosphate (NADPH). L-MB readily auto-oxidates to MB, a process which consumes molecular oxygen and yields hydrogen peroxide, a very strong oxidizing agent. The oxidizing conditions trigger the expression of a range of response genes, most notably the transcription factor Nrf2 (nuclear factor erythroid 2-related factor 2). Nrf2 interlinks with a range of cellular stress and homeostasis pathways, such as the oxidative and xenobiotic stress response, mitochondrial respiration, the UPR in the endoplasmic reticulum and autophagy (44,45). It is thus conceivable that the overexpression of Nrf2 has negative effects on SARS-CoV-2 infection. In support of this notion, Nrf2 mediated gene expression was found to be suppressed in biopsies obtained from COVID-19 patients (46). Intriguingly, another top performer in our screen, GPP-78 targets nicotinamide phosphoribosyltransferase (NAMPT), a crucial enzyme in the NAD salvage/recycling pathway (47).

An important and widely used clinical compound identified in our screen is MPA. MPA was previously reported to inhibit SARS-CoV-2 infection of humanized mice and human lung organoids (48). Given the immunosuppressive anti-inflammatory effects of MPA, we suggest a two-pronged modality of MPA towards treating COVID-19. One is an anti-viral action at moderate concentration to reduce systemic disease. MPA is a non-competitive inhibitor of ino-sine-5’-monophosphate dehydrogenase (IMPDH) involved in guanine biosynthesis (49). IMPDH interacts with and supports SARS-CoV nonstructural protein 14 (Nsp14), a guanine-N7-methyl-transferase implicated in virus replication and transcription (50). In addition, MPA leads to a reduction in the levels of the furin protease in human pluripotent stem cell derived lung organoids, and inhibits the entry of S-glycoprotein pseudotyped vesicular stomatitis particles (48). Furin has a key role in viral pathogenesis and dissemination, and is required for the limited proteolysis of the SARS-CoV-2 S-protein, and S-protein engagement with the neuropilin-1 receptor and virus infectivity, especially neuronal and heart cell infections (37,38).

The second modality of MPA has anti-inflammatory and immuno-suppressive effects, particularly at higher dosage of 10 µM plasma concentration, as demonstrated by the broad and long standing use of MPA and its prodrug formulation Mycophenolate Mofetil (MMF, Myfortic®, Novartis) in organ transplant patients, post operation for reduction of transplant rejection risk (51). Anti-inflammatory effects are considered to be beneficial to COVID-19 patients suffering from a dysregulated immune response, which includes a so-called cytokine storm (52). This has been shown with dexamethasone treatment of COVID-19 patients, either alone or in combination with remdesivir, albeit with variable outcomes and requiring close patient monitoring (53,54). In contrast, MPA / MFF has well characterized distinct anti-inflammatory effects. Reversible inhibition of IMPDH results in blockage of *de novo* guanosine synthesis, which suppresses the proliferation of T and B lymphocytes (55). B and T cells are largely devoid of salvage pathways for the synthesis of guanosine, and hence are particularly susceptible to MPA. MPA / MMF treatment leads to a reduction in mRNAs encoding pro-inflammatory cytokines, such as TNF-a, IL-6 &IL-1b (56,57). It further inhibits the activity of 6-pyruvoyl tetrahydro-biopterin synthase involved in the biosynthesis of tetrahydrobiopterin from GTP, a cofactor of aromatic amino acid monooxygenases and nitric oxide synthase. This inhibition leads to a reduction of the proinflammatory metabolite peroxynitrite (58), and could contribute to the beneficial effects of MPA / MFF in COVID-19. Along the same vein, SB-505124 was a triple hit against hCoV-229E-GFP, hCoV-OC43 and SARS-CoV-2 in our human cell culture screens. It is a reversible ATP competitive inhibitor of the TGF-beta type I receptor serine/threonine kinase, also known as activin receptor-like kinase (ALK) 4, as well as ALK5 and 7 (59). The TGF-beta receptor pathway controls differentiation, chemotaxis, proliferation, and activation of immune cells, and its inhibition by SB-505124 might contribute to a reduction in inflammation.

### Repurposing SARS-CoV-2 inhibitors in clinical development

One of our strongest hits in the cell culture experiments is MLN4924 exhibiting an excellent dose-response profile against hCoV-229E-GFP (EC_50_ 30 nM; TC_50_ 40.39 μM) and significant inhibition of SARS-CoV-2. MLN4924 (Pevonedistat) targets the Nedd8 activating enzyme (NAE) with potential anti-neoplastic activity, including cell growth inhibition through neddylation inhibition of Cullin ring ligases leading to ubiquitination failure, dysregulated cell cycle progression and apoptosis (60,61). It is a promising candidate in retinoblastoma therapy (62). MLN4924 also prevents the degradation of Nrf2, which has a central role in redox metabolism (63). Strikingly, this suggests that MLN4924 and MB might have similar mode-of-actions against SARS-CoV-2, namely dysregulating the oxidative state of the cell and activation of Nrf2.

Our best hit in cell cultures is Ro 48-8071 fumarate, a potent inhibitor of oxidosqualene cyclase (OSC) at an IC_50_ of 6.5 nM (64). OSC catalyzes the conversion of monooxidosqualene to lanosterol, and is in preclinical trials (64). Ro 48-8071 was originally developed to reduce plasma cholesterol levels. The sterol synthesis pathway plays a major role in membrane biology, and is involved in the replication process of many positive strand RNA viruses, including hepatitis C virus (65), poliovirus (66), rhinovirus (67) and SARS-CoV (68). Intriguingly, other high-quality hits, such as the azole-based antifungal drugs Posaconazole and Ravuconazole targeting sterol 14α demethylase (CYP51) (69,70) might block coronavirus infection by interfering with sterol biosynthesis. Among the original hits identified as effective against hCoV-229E, were two other azole-based antifungals, Mebendazole (discarded because of autofluorescence in follow-up experiments) and Isavuconazole. This finding further highlights the significance of the cholesterol biosynthesis pathway and its potential as a drug target against SARS-CoV-2. Posaconazole has been in clinical use since 2006 and generics thereof are available, making it an interesting candidate for transition to clincal trials.

### Other compounds

Our screen reveals several additional launched compounds already described for their potential to treat COVID-19, including Fostamatinib, Eprinomectin and Betulinic acid. Fostamatinib is an FDA approved Syk tyrosine kinase inhibitor for the treatment of chronic immune thrombocytopenia (71), and is in a phase 3 clinical trial against COVID-19 (<u>NCT04629703</u>). Although Fostamatinib inhibited SARS-CoV-2 in our cultured human cells and African green monkey Vero cells, we did not pursue this compound further due to morphological alterations in the HAEEC, including out-growth of fibroblasts upon prolonged exposure to Fostamatinib. Likewise, we did not follow up on Eprinomectin, a benzodiazepine receptor agonist with anti-parasitic activity but unknown mode-of-action, though approved for veterinary use in horses, cattle and cats, akin to other avermectins (72). We did not further investigate Betulinic acid, a pentacyclic triterpenoid with attributed anti-retroviral, anti-malarial, and anti-inflammatory properties but also anti-proliferative and apoptotic effects (73).

The screen further reveals a cluster of five distinct compounds targeting glycogen synthase kinase 3 (GSK-3), namely LY2090314 (74), GSK-3 inhibitor IX (75), TCS-21311 (76), AZD2858 (77) and CHIR-98014 (78). Remarkably, LY2090314 was the most potent compound identified in our hCoV-229E-GFP screen with an EC_50_ of 2 nM. GSK-3 phosphorylates a variety of proteins in glycogen metabolism, innate immunity and apoptosis (79). Different GSK-3 inhibitors have been approved to treat cancer, inflammation, Alzheimer’s, and diabetes (80), and were proposed for SARS-CoV-2 treatment. The latter is based on the notion that GSK-3 is implicated in phosphorylation of SARS-CoV-2 N-protein (81). Unfortunately, none of the GSK-3 inhibitors identified in our screen translated anti-viral effects to SARS-CoV-2 infections.

In conclusion, our image-based, high content, full infection cycle screen against human coronavirus infection provides a high quality hit list of preclinical and clinical compounds with high repurposing potential. We identified five clinically launched compounds. Two of these were tested in and found to inhibit virus egress from HAEEC (MB and MPA). Posaconazole, Thonzonium bromide and Cetylpyridinium blocked virus dissemination in 2D cell cultures. Except for Cetylpyridinium, all of these clinically launched compounds are orally administered.

## Materials &Methods

### Viruses

hCoV-229E-GFP and SARS-CoV-2 (TAR clone 3.3, München-1.1/2020/929) were obtained from Dr. Volker Thiel (University of Bern) (82,83). Human CoV-229E-GFP was plaque purified and expanded on Huh7 cells for 48h. Supernatant was collected and cleared by centrifugation at 5000 x g for 10 min. Human CoV-OC43 (ATCC-VR-1558) was obtained from LGC Standards Limited, (Teddington, United Kingdom). and expanded as described above. Virus titers were determined by FFU titration according to GFP or immunofluorescence signal, and by TCID_50_ titration according to the Spearman-Kärber method.

### Cell lines

Huh7 and VeroE6 cells were obtained from Dr. Volker Thiel (University of Bern, Switzerland). Huh7-ACE2, HeLa-ACE2 and A549-ACE2 were generated by stable transfection with a lentiviral vector (pLVX-ACE2-IRES-BSD). Parental HeLa and A549 cells were obtained from ATCC. The parental HeLa cell line additionally expressed an inducible eCas9 and a non-targeting sgRNA.

### Cell culture

Huh7, Huh7-ACE2, HeLa-ACE2 and A549-ACE2 cells were maintained in Dulbecco’s Modified Eagle Medium (D6429; Sigma-Aldrich, St. Louis, USA) supplemented with 10% fetal bovine serum (FBS, 10270; Invitrogen, Carlsbad, USA), Non-essential amino acids (M7145; Sigma-Aldrich, St. Louis, USA) and subcultured by PBS washing and trypsinisation (C-41020; Trypsin-EDTA, Sigma-Aldrich, St. Louis, USA) twice weekly. Cell lines were kept at 37 °C, 5% CO2, 95% humidity. Infections with Huh7 cells were conducted with Minimal Essential Medium (MEM, Sigma-Alrich, St. Louis, USA) supplemented with 10% FBS.

### Bronchial and nasal HAEEC

Human nasal and bronchial airway tissue (MucilAir™, Epithelix Sarl, Geneva, Switzerland) cultured on transwell inserts (24-well plate) were maintained at air-liquid interface according to the supplier’s instructions and cell culture medium (EP05MM). The bronchial HAEEC were obtained from individual donors (Batch nr: MD0810) and the nasal HAEEC from a pool of fourteen donors (Batch nr: MP0009). All donors were non-smoker and healthy.

### SARS-CoV2 infection of the MucilAir™ tissue and drug treatment

Three days prior infection, MucilAir™ tissue apical surface were washed with 200 µl of warmed MucilAir™ culture medium for 20 minutes at 37°C to homogenize the amount of mucus between inserts. Inserts were inoculated on the apical side with 1,000 TCID_50_ of SARS-CoV2 in a final volume of 100 µl at 37°C for 2 h. Then the SARS-CoV2 inoculum was removed and the apical surfaces quickly washed two-times with PBS. SARS-CoV2 produced at the apical surface were collected at different time post infection by 20 min apical wash and quantified in parallel by rt-qPCR and TCID_50_ titration in VeroE6 cells. Drug treatment were done at the indicated times pre and/or post infection by addition of the indicated drug concentrations in fresh basolateral medium. Nasal and bronchial MucilAir™ tissue analysis were performed in duplicate. Statistical details of experiments can be found in the figure legends. Statistical analyses were performed using GraphPad Prism 8.

### SARS-CoV2 RNA quantification by real-time RT-qPCR

SARS-CoV2 RNAs were extracted from samples with the Quick-RNA™ MiniprepPlus Kit (Zymo, R1058) after TRIzol™ reagent (Invitrogen, 15596026) treatment, according to the manufacturer’s instructions. The number of viral genome copies was evaluated by one-step real-time RT-qPCR of the SARS-CoV2 M-gene with the Superscript TMIII Platinum One step Quantitative Kit (Invitrogen, 11732-020). Five µl of extracted RNAs were added to 15 µl of reaction mixture containing: reaction mix buffer (1x), 0.5 µl of MgSO_4_ buffer, reference dye ROX (50 nM), each primer (400 nM), Taqman probe (100 nM), and 0.4 µl SuperScript III RT/Platinum® TaqMix. The final volume was made up to 20 µl with nuclease-free water. Thermal cycling was performed in a QuantStudio 3 Real-Time PCR System thermocycler (Thermo Fisher Scientific) in MicroAmp® Optical 96-well reaction plates (Applied Biosystems, N8010560).

Conditions were: reverse transcription for 20 minutes at 50°C, Taq DNA polymerase activation for 2 minutes at 95°C, and then 45 cycles of amplification consisting of DNA denaturation for 15 s at 95°C, and combined annealing/extension for 1 minute at 60°C. Fluorescence data were collected at the end of each cycle. The number of viral genome copies was evaluated from a standard curve amplified in parallel of 10-fold serially diluted *in vitro* transcribed SARS-CoV2 M-gene RNAs quantitative standards (SARS-qRNAs) in RNase-free water. The SARS-qRNAs were *in vitro* transcribed from a synthetic SARS-CoV2 M-gene DNA template (Microsynth AG, Balgach Switzerland) under control of a T7 promoter by using the HiScribe™ T7 High Yield RNA Synthesis Kit (NEB, E2040). Then synthesized SARS-qRNAs were quantified with a NanoDrop spectrophotometer (Thermo Fisher Scientific). The oligonucleotides are presented in Table S3.

### Compound library

The Prestwick chemical library (PCL) was purchased from Prestwick Chemical (Illkirch, France). The remaining compounds were purchased from abcr GmbH, AK Scientific, Inc., Abcam plc., Acros Organics, AdipoGen Life Sciences, Inc., Advanced ChemBlocks Inc., Alinda Chemical Trade Company Ltd., Angene, Apollo Scientific Ltd., Axon Medchem, BIOTREND Chemikalien GmbH, BLD Pharmatech Ltd., Biosynth Carbosynth, Cayman Chemical Company, ChemBridge Corporation, ChemDiv Inc., Chemodex Ltd., Cohesion Biosciences Ltd., Combi-Blocks, Inc., Enamine Ltd., Enzo Life Sciences Inc., Fluorochem Ltd., Focus Biomolecules, InterBioScreen Ltd., J&K Scientific Ltd., Key Organics Ltd., LabSeeker, Inc, Life Chemicals Inc., Matrix Scientific, Maybridge Ltd., MedChemExpress, Merck KGaA, Otava Ltd., Pharmeks Ltd., Ramidus AB, SYN-kinase, Santa Cruz Biotechnology, Selleck Chemicals LLC, Sigma-Aldrich, Specs, Target Molecule Corp., Tocris Bioscience, Tokyo Chemical Industry, Toronto Research Chaemicals, UkrOrgSyntez Ltd., Vitas-M Laboratory Ltd. or Wuhan ChemFaces Biochemical Co., Ltd.. Compounds were dissolved in DMSO at a concentration of 10 mM.

### High throughput compound screening

The screen was conducted in 4 completely independent biological replicates without technical replicates. For fluidics handling, a Matrix WellMate dispenser and Matrix WellMate tubing cartridges (Thermo Fisher Scientific, Waltham, USA), an Assist Plus pipetting robot (Integra Biosciences AG, Zizers, Switzerland) and an Echo acoustic dispenser (Labcyte Inc., Indiana, USA) were used. Compounds were spotted at 1.67 μM final concentration in microscopy grade 384-well plates (Perkin-Elmer, Waltham, USA), and frozen. Pre-spotted plates were thawed at room temperature 30 minutes prior to cell seeding. 6000 Huh7 cells were seeded in 25 μL/well MEM +10% FCS and incubated at 37°C, 5% CO_2_, 95% humidity overnight. Infection was performed with 50 focus forming units (FFU) per well in 5 μL and incubated at 33.5°C, 5% CO2, 95% humidity for 48h. Finally, cells were fixed by adding 10 μL 16% paraformaldehyde (Sigma-Aldrich, St. Louis, USA) containing 0.4 μg/mL Hoechst 33342 (Sigma-Aldrich, St. Louis, USA). The fixation reaction was quenched in 25 mM NH_4_Cl in phosphate-buffered saline (PBS) for 30 minutes. Plates were washed three times with PBS. Finally, PBS was replaced with PBS + 0.02% N_3_. Plates were imaged with an automated inverted epifluorescence microscope at 4x magnification (Molecular Devices, San Jose, USA). For the validation experiments with hCoV-OC43 and SARS-CoV-2, compounds were spotted at 1.75 μM, cells seeded in 80 μL, incubated at 37°C, 5% CO2 and 95% humidity overnight and infected with 120 FFU in 20 μL. After 68 hours at 33.5°C (OC43) or 24 h or 48 h at 37°C (SARS-CoV-2), cells were fixed as described above. Subsequently, the experiments were subjected to immunostaining. Cells were permeabilized with 0.2% Triton-X-100 for 5 minutes, washed with PBS, incubated with primary antibodies against coronavirus nucleoprotein (Chemicon MAB 9013 for OC43, Rockland 200-401-A50 for SARS-CoV-2) in PBS + 1% bovine serum albumin (BSA) for 1 h, washed and incubated with a secondary antibody in PBS + 1% BSA for 1h. Cells were then washed again, PBS was replaced with PBS + 0.02% N_3_ and finally, plates were imaged as described above.

### Image analysis

Viral infection and cell health was parametrized using Plaque2.0 (18). The read-out includes the 5 main parameters: number of nuclei, number of infected nuclei, infection index, total virus intensity and number of plaques/infection foci.

### Post-processing

Results obtained by image analysis with Plaque2.0 were annotated and filtered using R version 4.0.2 in RStudio Version 1.3.1056. Readout values were per-plate normalized by the mean values of the DMSO controls. Compounds were considered toxic if the mean number of nuclei was lower than the mean number of nuclei of all negative control wells minus 2 times the standard deviation of the number of nuclei of all negative control wells. Compounds were considered hits if the mean value of all replicates falls below the mean of all negative control wells minus 3 times the standard deviation of all negative control wells for that specific readout. Hit lists were combined for all parameters. In the validation experiments, compounds were considered toxic if the mean number of nuclei was lower than the mean number of nuclei of all negative control wells divided by 2.

## Author contributions

UFG initiated, conceived and supervised the project. LM, RV, NM, LY, DS, AGG, AP, DO, MB, MS, FK, GT, UFG designed, carried out or analyzed experiments. LM, RV provided data visualization. VA, LM provided data curation and software. LM, RV, UFG wrote the paper. GT, UFG acquired funding. All authors read and approved the manuscript.

The authors declare no competing interests.

## Acknowledgements

We acknowledge financial support from the Swiss National Science foundation (SNSF) to UFG and GT (31CA30_196177 / 1), the NCCR Chemical Biology supported by the SNSF for purchasing the BSF-EPFL repurposing collection, and a special Coronavirus Research grant from the University of Zurich to UFG. We are grateful to Dr. Adriano Aguzzi and Dr. Simone Hornemann for granting generous access to their BSL3 laboratory. We thank Dr. Volker Thiel and Dr. Nadine Ebert for their kind gifts of hCoV-229E-GFP, SARS-CoV-2 isolate Munich, cell lines, as well as discussions and hands-on information, as well as Dr. Daria Seiler, Dr. Fanny Georgi, Jonathan Vesin, Antoine Gibelin and Julien Bortoli for their technical assistance.

