## Supplementary figures and tables for "Arrayed multicycle drug screens identify broadly acting chemical inhibitors for repurposing against SARS-CoV-2"

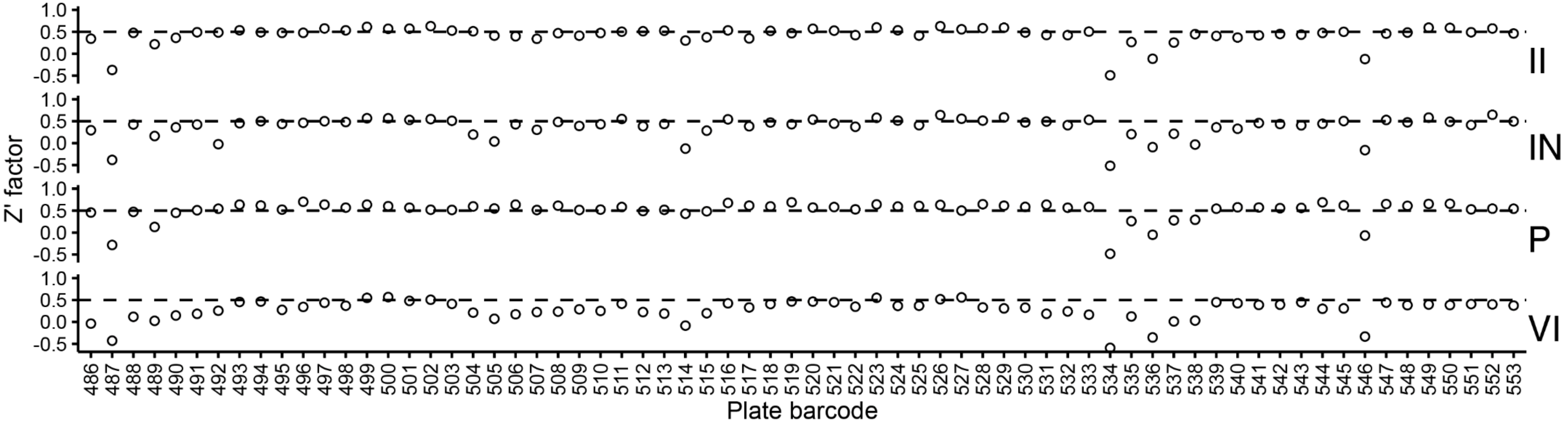


**Supplementary figure 1: Z' factors of each screening plate.** Each facet depicts the per-plate Z' factors for the assessed parameters. II: infection index, IN: infected nuclei, P: plaques, VI: total virus intensity. The dashed line intersects the y-axis at 0.5. Mean Z' factors are II = 0.432, IN = 0.378, P = 0.509, VI = 0.276.

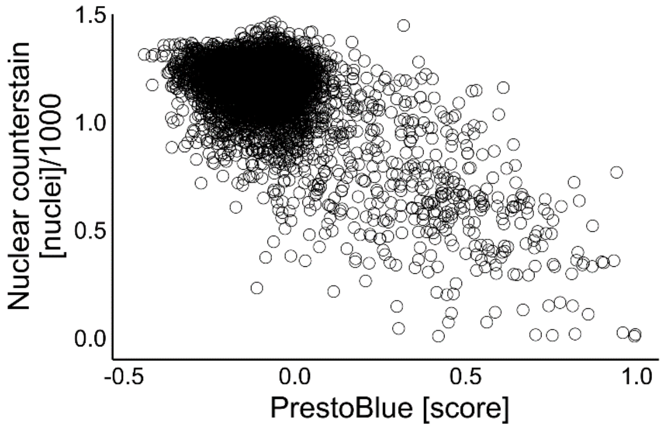


**Supplementary figure 2: Comparison between nuclear counterstain and PrestoBlue as a measurement for cell viability.** The number of nuclei was assessed based on the nuclear counterstain in the screening setup. In contrast, the PrestoBlue based assay was performed in absence of virus, but all the other parameters were kept the same. The score of the Prestoblue assay was normalized with the positive control set as 1, and the negative control as 0. As a result, there is a negative correlation between the parameters.

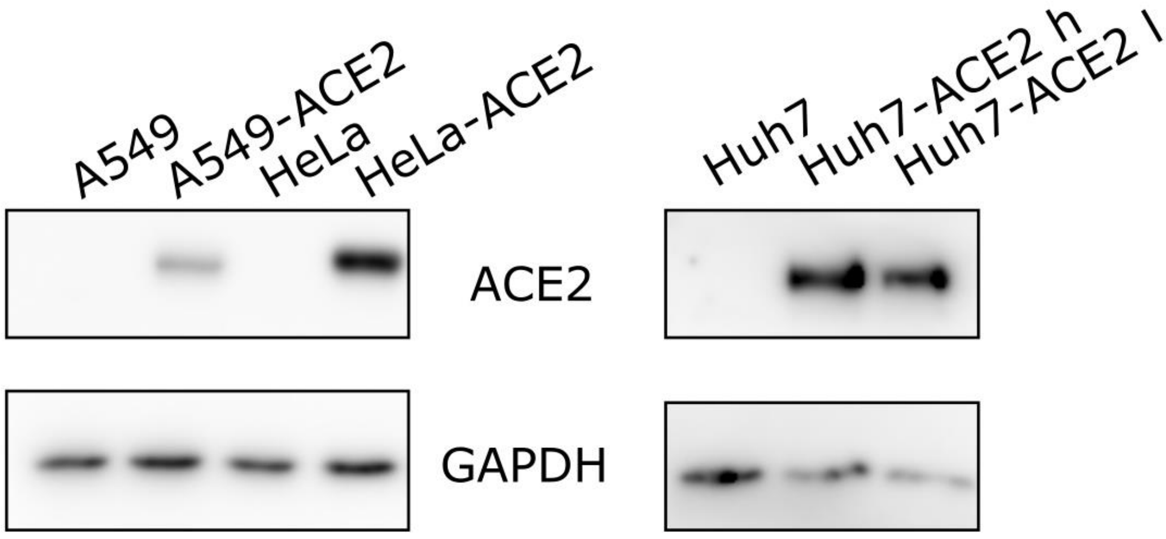


**Supplementary figure 3: ACE2 expression of lentivirus transduced cell lines.** ACE2 expression was verified in all cell lines transduced with lentivirus encoding human ACE2. An unspecific band was observed in both the untransduced and the transduced HeLa cells (not shown). The GAPDH loading control for the left blot including A549 and HeLa cell lines was performed with the same lysates, but on separate lanes. The two Huh7-ACE2 lanes come from separate cell populations, transduced with either a high or a low MOI of lentivirus. For the present work, we used the cell line on the right (low MOI transduction) because ACE2 expression was sufficient.

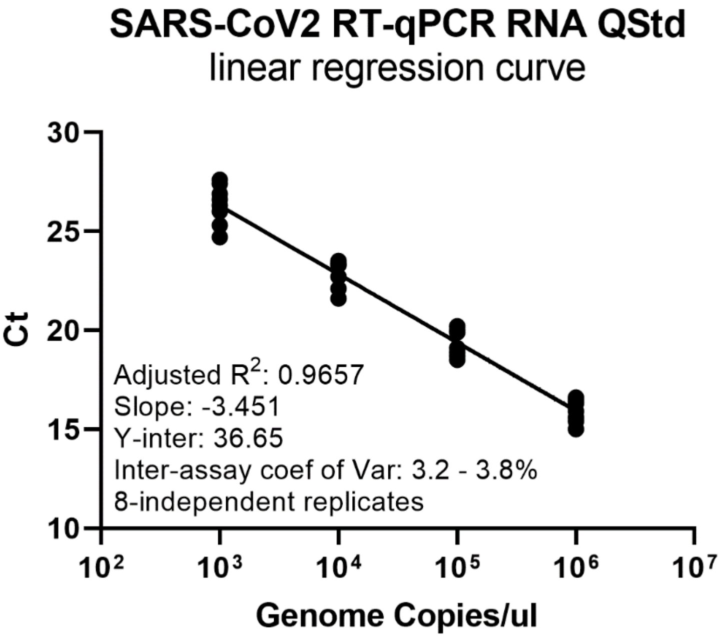


**Supplementary figure 4. Inter-assay reproducibility of the SARS-CoV2 membrane protein gene (M-gene) quantitative RT-PCR.** The linear regression curve was generated by plotting the Ct values against the copy numbers (106 to 103 copies per µl) of the RNA M-gene quantitative standards in 8-independent replicates. The adjusted coefficient of correlation (R2) and the inter-assay coefficient of correlation are represented. The coefficient of correlation was calculated for each quantitative point according the formula: %CV= (SD / Ct mean)×100.

| Compound | Original indication | Clinical status | $EC_{50}$ [µM] | $TC_{50}$ [µM] | Susceptible cell lines (n.) | Administration |
|---|---|---|---|---|---|---|
| A0001 | Friedreich ataxia | Phase II | 0.50 | 8.40 | 0 | Systemic |
| Abafungin | Antifungal | Phase III | 1.64 | 4.67 | 1 | Topical |
| Acivicin | Cancer | Phase I | 0.67 | >50 | 0 | Systemic |
| Adarotene | Antibiotic | Phase I | 1.29 | 1.84 | 0 | Systemic |
| Amuvatinib | Cancer | Phase II | 0.94 | 9.21 | 0 | Systemic |
| Anisomycin | Antibiotic | Phase I | 0.14 | 11.10 | 0 | Topical |
| APY0201 | Immunosuppressant | Preclinical | 3.36 | 21.19 | 1 | Systemic |
| AT9283 | Cancer | Phase II | 0.45 | 21.29 | 0 | Systemic |
| AZD-2858 | Bone fracture | Phase I | 0.39 | 6.39 | 0 | Systemic |
| AZD-5438 | Cancer | Preclinical | 1.08 | 13.99 | 1 | Systemic |
| Benzyldimethyl hexadecylammonium | Antiseptic | Preclinical | 0.73 | 3.49 | 1 | Topical |
| Betulinic acid | Cancer | Phase II | 0.18 | 18.06 | 0 | Topical |
| Cerdulatinib | Cancer | Phase II | 1.12 | 11.29 | 0 | Systemic |
| Cetylpyridinium | Antiseptic | Launched | 0.24 | 1.62 | 2 | Topical |
| CHIR-124 | Cancer | Preclinical | 0.51 | 11.33 | 1 | Systemic |
| CHIR-98014 | Cancer | Preclinical | 0.26 | >50 | 0 | Systemic |
| CUDC-907 | Cancer | Phase II | 0.03 | 0.03 | 0 | Systemic |
| Cyclopiazonic acid | Diabetes | Preclinical | 0.87 | 12.65 | 0 | Topical |
| Eprinomectin | Vet. Antiparasitic | Veterinary | 0.59 | 4.19 | 0 | Topical |
| FH535 | Cancer | Preclinical | 0.74 | 2.54 | 0 | Systemic |
| Geldanamycin | Antibiotic | Phase II | 0.03 | 0.02 | 0 | Systemic |
| GPP-78 | Spinal cord injury | Preclinical | 0.84 | 10.12 | 2 | Systemic |
| GSK 3 Inhibitor IX | Cancer | Phase II | 0.46 | 8.80 | 1 | Systemic |
| GZD824 | Cancer | Phase II | 0.49 | 0.98 | 1 | Systemic |
| Isavuconazole | Antifungal | Phase III | 0.49 | 7.15 | 0 | Systemic |
| JTE-013 | Immunosuppressant | Preclinical | 1.26 | 10.10 | 0 | Systemic |
| LE-135 | Cancer | Preclinical | 0.38 | 4.98 | 0 | Systemic |
| Lomefloxacin HCl | Antibiotic | Launched | | | 0 | Topical |
| LY2090314 | Cancer | Phase II | 0.00 | 31.66 | 1 | Systemic |
| Mebendazole | Anthelmitic | Launched | 46.23 | 37.22 | 1 | Topical |
| Methylene blue | Methemoglobinemia | Launched | 1.43 | 8.71 | 2 | Systemic |
| MLN4924 | Cancer | Phase II | 0.03 | 40.39 | 3 | Systemic |
| Mycophenolic acid | Immunosuppressant | Launched | 1.85 | 45.82 | 3 | Systemic |
| OTSSP167 | Cancer | Phase I | 0.04 | 0.77 | 0 | Systemic |
| Oxaprozin | Immunosuppressant | Launched | | | 0 | Systemic |
| Pelitinib | Cancer | Phase II | 1.95 | 8.48 | 1 | Systemic |
| PF-04457845 | Osteoarthritis | Phase II | 1.18 | 6.09 | 0 | Systemic |
| PF-3845 | Antinociceptive | Preclinical | 1.04 | 5.78 | 0 | Systemic |
| PFK-015 | Cancer | Phase I | 1.97 | 3.50 | 0 | Systemic |
| Posaconazole | Antifungal | Launched | 4.20 | >50 | 4 | Systemic |
| Fostamatinib | Immune thrombo-cytopenic purpura | Launched | 1.33 | 14.71 | 2 | Systemic |
| RAF265 | Cancer | Phase II | 0.70 | 6.96 | 1 | Systemic |
| Ravuconazole | Antifungal | Phase II | 0.86 | 5.95 | 2 | Systemic |
| Ro 48-8071 | Cancer, Antiviral | Preclinical | 1.86 | 8.64 | 4 | Systemic |
| SB225002 | Cancer | Preclinical | 2.31 | 9.45 | 2 | Systemic |
| SB505124 | Cancer | Preclinical | 7.48 | 22.12 | 0 | Systemic |
| Tanaproget | Contraceptive | Phase II | 0.48 | 5.47 | 0 | Systemic |
| TCS-21311 | Immunosuppressant | Preclinical | 0.19 | 19.07 | 1 | Systemic |
| Thonzonium bromide | Antiseptic | Launched | 0.78 | 4.59 | 3 | Topical |
| TRO 19622 | Spinal muscular atrophy | Phase II | | | 0 | Systemic |
| UK-5099 | Cancer | Preclinical | 0.35 | 20.64 | 0 | Systemic |
| Verteporfin | Wet macular degen. | Launched | 0.87 | 2.07 | 1 | Systemic |
| VU29 | mGlu5R modulator | Preclinical | 8.14 | 39.42 | 0 | Systemic |

**Supplementary table 1. Overview of all hits.**

| Drug (replicate)[a] | Apical 1-dpi (RT-qPCR, Ct-value)[b] M-gene | Pol-gene | ΔCt |
|---|---|---|---|
| **Pre-treatment (10uM)** | | | |
| Cpd 16 (A) | 29.4 | 28.3 | 1.1 |
| Cpd 16 (B) | 29.9 | 29.0 | 0.9 |
| Cpd 17 (A) | 33.8 | 33.4 | 0.4 |
| Cpd 17 (B) | 35.5 | 32.9 | 2.6 |
| Cpd 31 (A) | 30.0 | 29.0 | 1.0 |
| Cpd 31 (B) | 30.1 | 28.4 | 1.7 |
| Remdesivir (A) | 33.2 | 32.4 | 0.8 |
| Remdesivir (B) | 31.9 | 31.8 | 0.1 |
| **Pre-treatment (1uM)** | | | |
| Cpd 16 (A) | 32.6 | 40.1 | 7.5 |
| Cpd 16 (B) | 30.1 | 28.7 | 1.4 |
| Cpd 17 (A) | 33.3 | 32.2 | 1.1 |
| Cpd 17 (B) | 31.9 | 31.3 | 0.6 |
| Cpd 31 (A) | 28.6 | 27.9 | 0.7 |
| Cpd 31 (B) | 30.2 | 29.9 | 0.3 |
| **Post-treatment (10uM)** | | | |
| Cpd 16 (A) | 31.0 | 29.4 | 1.6 |
| Cpd 16 (B) | 29.3 | 28.8 | 0.5 |
| Cpd 17 (A) | 30.9 | 29.9 | 1.0 |
| Cpd 17 (B) | 27.8 | 28.1 | 0.3 |
| Cpd 31 (A) | 30.2 | 29.2 | 1.0 |
| Cpd 31 (B) | 26.0 | 26.4 | 0.4 |
| Remdesivir (A) | 27.3 | 26.0 | 1.3 |
| Remdesivir (B) | 25.7 | 25.4 | 0.3 |
| **Post-treatment (1uM)** | | | |
| Cpd 16 (A) | 31.0 | 29.4 | 1.6 |
| Cpd 16 (B) | 29.2 | 28.0 | 1.2 |
| Cpd 17 (A) | 30.5 | 29.4 | 1.1 |
| Cpd 17 (B) | 28.1 | 28.2 | 0.1 |
| Cpd 31 (A) | 29.7 | 27.9 | 1.8 |
| Cpd 31 (B) | 30.4 | 32.8 | 2.4 |
| **Control** | | | |
| DMSO (A) | 34.5 | 32.8 | 1.7 |
| DMSO (B) | 31.2 | 29.9 | 1.3 |

[a] Nasal MucilAir from December 2020

[b] Fluorescence threshold at 0.025 ΔRn

**Supplementary table 2. SARS-CoV-2 M and nsp12 gene RT-qPCR threshold cycle comparison.**

| Name | DNA Sequence (5’ to 3’) |
|---|---|
| SARS2-Mgene-FP | TTGGATCACCGGTGGAATTG |
| SARS2-Mgene-RP | GCAATGAAGTAGCTGAGCCAC |
| SARS2-Mgene-Pb [a] | *HEX*-TCGCAATGGCTTGTCTTGTAGGCTTG-*BHQ1* |
| SARS2-Mgene-QStd [b] | GGAAGCTT**TAATACGACTCACTATAGGG**CTGTTTACAGAATAAATTGGATCACCGGTGGAATTGCTATCGCAATGGCTTGTCTTGTAGGCTTGATGTGGCTCAGCTACTTCATTGCTTCTTTCAGACTGTTTGCGCGTACGCGTTCCATG |
| SARS2-nsp12-FP | CTCACCTTATGGGTTGGGAT |
| SARS2-nsp12-RP | CACGTTGTATGTTTGCGAGC |
| SARS2-nsp12-Pb [a] | *FAM*-GTGATAGAGCCATGCCTAACATGC-*BHQ1* |

**Supplementary table 3.**

[a] The Taqman probes (Pb) are labelled in 5’ with the hexachloro-fluorescein (HEX) or 6-Carboxyfluorescein (FAM) and in 3’ with the black hole quencher-1 (BHQ1). Both are indicated in italic.

[b] The T7 promoter sequence used for the RNA *in vitro* transcription is indicated in bold. The oligos hybridization sequences are underlined. Abbreviations: forward primer (FP), reverse primer (RP), TaqMan probe (Pb), quantitative standard (Qstd)
